# patcHwork: A user-friendly pH sensitivity analysis web server for protein sequences and structures

**DOI:** 10.1101/2022.02.02.478804

**Authors:** Mirko Schmitz, Anne Schultze, Raimonds Vanags, Karsten Voigt, Barbara Di Ventura, Mehmet Ali Öztürk

## Abstract

pH regulates protein function and interactions by altering the charge of individual residues causing the loss or gain of intra-molecular non-covalent bonds, which may additionally lead to structural rearrangements. While tools to analyze residue-specific charge distribution of protein sequences and structures at a given pH exist, currently no tool is available to investigate non-covalent bond changes at two different pH values. In an effort to make protein pH sensitivity analysis more accessible to researchers without computational structural biology background, we developed patcHwork, a web server that combines the identification of amino acids undergoing a charge shift with the determination of affected non-covalent bonds at two user-defined pH values. At the sequence-only level, patcHwork applies the Henderson-Hasselbalch equation to determine pH-sensitive residues. When the 3D protein structure is available, patcHwork can be employed to gain a deeper mechanistic understanding of the effect of pH on a protein of interest. This is achieved using the PDB2PQR and PROPKA tools and non-covalent bond determination algorithms. A user-friendly interface allows visualizing pH-sensitive residues as well as affected salt bridges, hydrogen bonds and aromatic (pi-pi and cation-pi) interactions. Importantly, patcHwork can be used to identify patches, a new concept we propose of pH-sensitive residues in close proximity on the protein structure, which may have a major impact on function. We demonstrate the attractiveness of patcHwork studying experimentally investigated pH-sensitive proteins. (Access:https://patchwork.biologie.uni-freiburg.de/)

**Graphical abstract:** 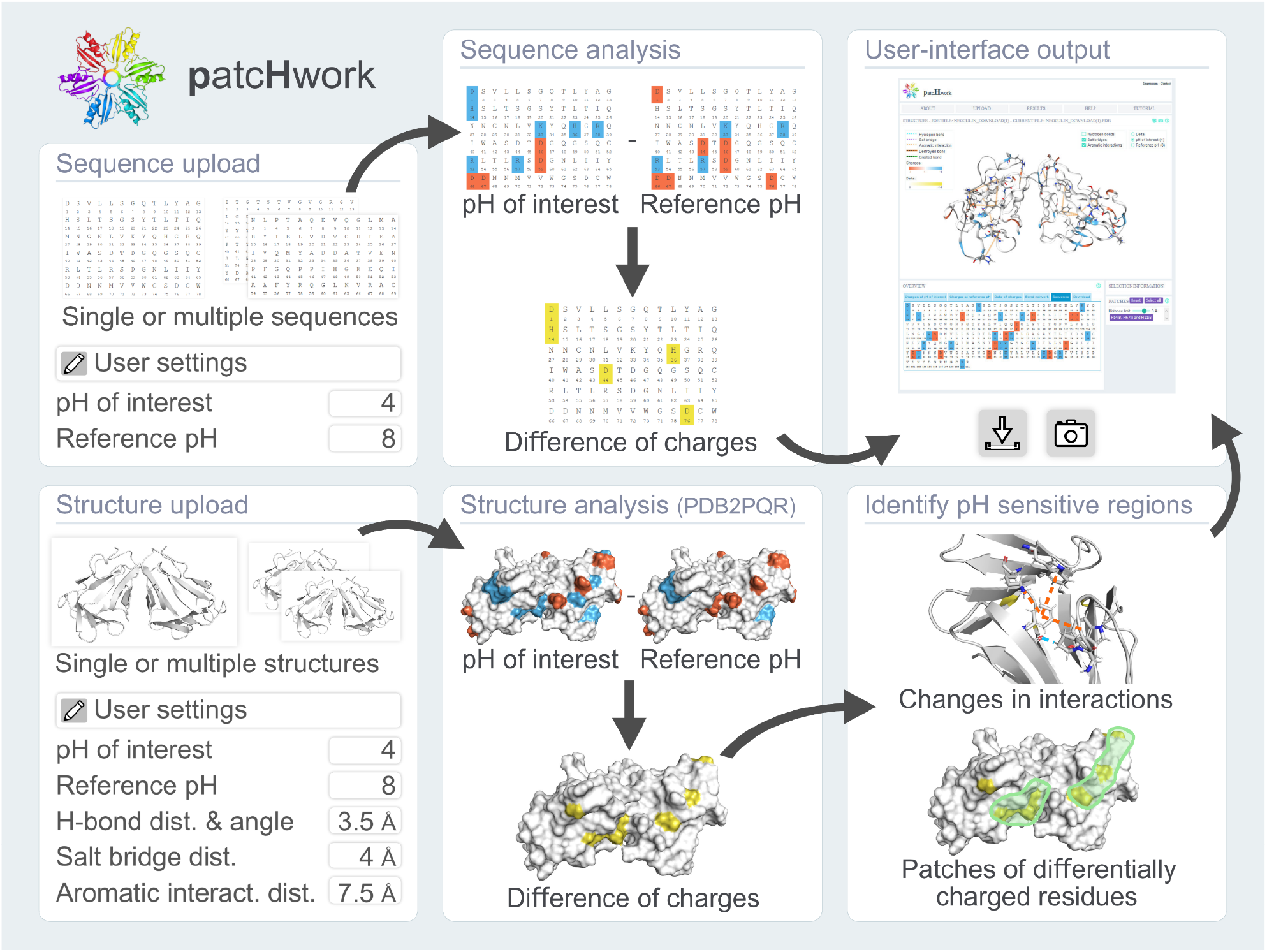

## Introduction

The concentration of hydrogen ions in a solution (used to calculate the pH) determines the charge of the side chains of the amino acids in a given protein by regulating their protonation state. Various properties such as protein solubility (1), stability (2), ability to interact with other molecules (3), flexibility (4) and activity (5) are affected by pH. As a matter of fact, the amount of protonation of the amino acids’ side chains in proteins has been proposed to be a new form of protein post-translational modification (6). Ionizable residues in proteins react to the surrounding pH according to their ionization constant (pKa) and they become positively or negatively charged in pH values below or above their pKa, respectively. This adaptation can cause loss or gain of intra-molecular non-covalent bonds between residues, which can subsequently result in structural rearrangements that could regulate protein function and interactions (7). Thus, it is crucial to understand the effect of pH change on a protein structure and determine any resulting adjustments in intra-molecular non-covalent bonds.

Currently, a number of web servers exist that allow calculating the charges of the amino acid side chains at a certain pH using either sequence or structural information (8–10).VOLPES (8) for instance applies the Henderson-Hasselbalch equation (11) to calculate the charges of the side chains of the amino acids in a protein of interest at a user-defined pH using only sequence information. However, the Henderson-Hasselbalch equation does not account for the influence of neighboring amino acids on the pKa of a residue side chain (12). An analysis at the level of the structure is, therefore, clearly required to precisely determine the effect of pH change on proteins.

Structure-based analysis of pH-mediated changes in the charge of the side chains of amino acids has benefited from datasets obtained from NMR experiments, which significantly helped to refine computational methods for pKa prediction of residues in protein structures (13). For example APBS (10) calculates the electrostatic potentials of proteins by assigning charge and radius to atoms using the PDB2PQR (14) and PROPKA (15) tools. Protein-sol (8) provides overall pH-dependent charge information for proteins of interest, as well as predictions of the destabilization of residue-specific electrostatic interactions due to limited ionization ability of buried amino acids (12). There is also a graphical user interface (GUI) plug-in implementation (16) on the VMD software (17) to visualize PROPKA (15) predictions. While being useful, these approaches do not allow the user to easily monitor the appearance / disappearance of intra-molecular non-covalent bonds (salt bridges, hydrogen bonds, pi-pi and cation-pi interactions) when the pH is changed from one value to another. On the other hand, these bonds can be investigated only at the default physiological pH with web servers such as RING 2.0 (18), Arpeggio (19) and ProteinTools (20). Taken together, currently no tool offers the possibility to directly observe protonation changes of amino acids caused by a shift in pH between two user-defined values and the resulting gain/loss of non-covalent bonds in the protein structure. Therefore, researchers wishing to analyze pH sensitivity of a given protein need to use several tools in parallel with manual curation of the outputs, which makes such an analysis difficult for researchers with limited computational structural biology knowledge.

Here we present patcHwork, a novel web server that supports high-throughput pH sensitivity analysis at two user-defined pH values either at the sequence or structure level. At the sequence level, patcHwork allows submitting up to ten thousand protein sequences, which are then analyzed using the Henderson-Hasselbalch equation (11) to determine pH-sensitive residues at the user-defined pH values. When the 3D protein structure is available, further mechanistic understanding of the effect of pH on a protein of interest can be obtained by the execution of the PDB2PQR (14) and PROPKA (15) software and non-covalent bond determination algorithms (21–24). pH-sensitive residues and pH-mediated changes in salt bridges, hydrogen bonds, and aromatic (pi-pi and cation-pi) interactions are visualized in an interactive GUI. Additionally, users obtain information regarding so-called patches, groups of pH-sensitive residues found in a customizable physical distance on the protein structure, which may play a more profound role than individual amino acids in the regulation of protein function upon pH change. To demonstrate the workflow and the power of patcHwork, we carried out the sequence-based pH sensitivity analysis of *E. coli* cell envelope proteins, structures of the taste-modifying protein neoculin (25, 26) and the pH-regulated mouse anion exchanger 2 (AE2) protein (27–31).

## Functionalities of patcHwork

patcHwork has four main computational components to investigate the response of proteins to a change in pH: protein sequence-based analysis, protein structure-based analysis, non-covalent bond analysis, and identification of pH-sensitive patches.

### Protein sequence-based analysis

The Henderson-Hasselbalch equation (11) is solved for each amino acid of the submitted protein FASTA sequences for two user-given pH values. Residue-specific charges for each pH value and also delta charges are provided as an interactive output. In order to rank the proteins based on their pH sensitivity, an “overall charge score” is defined as follows: for each protein, the sum of the charges at the pH of interest is subtracted by the sum of charges at the reference pH and then normalized by the total number of residues in the protein.

### Protein structure-based and non-covalent bond analyses

The protonation state of each amino acid in the submitted protein PDB structures is calculated using the PDB2PQR (14) and PROPKA (15) tools for the two user-given pH values. In addition to residue-specific charge information, created and destroyed non-covalent bonds (salt bridges, hydrogen bonds and pi-pi and cation-pi interactions) upon pH change are determined and given as an interactive output.

### pH-sensitive patches

On protein structures, residues that change their protonation state at a given pH shift and are found within a radius of ≤ 8 Å from each other (customizable) are defined as a pH-sensitive patch.

Further details of sequence and structure-based analyses, pH-sensitive patch identification and non-covalent bond determination, as well as the web server implementation are given in the Supplementary Information.

## Case studies

### Sequence-based pH sensitivity analysis of *E. coli* cell envelope proteins

To demonstrate the advantage of using patcHwork for pH sensitivity analysis, we asked whether we could identify in a high-throughput manner proteins that are mostly affected by pH looking at a family of proteins that is exposed to the extracellular medium and, consequently, is most likely affected by pH change than cytoplasmic proteins: cell envelope proteins. We collected the FASTA sequences of 309 *E. coli* proteins annotated as being part of the cell envelope (GO:0030313) and determined for each the overall charge score (residues’ total charge shifts normalized by the protein length) when changing the pH from 1 to 14 with an increment of 1 (for details see Supplementary Information). Next we created for each protein a delta score (Δ) taking the mean of the overall charge scores of all pH shifts (Figure 1). We propose that ranking the proteins based on the Δ score we are able to identify the most and least pH-responsive proteins, respectively. After obtaining this ranked list, we checked the literature for experimental information on the top and bottom five proteins to verify whether they are reported to be involved in pH regulation / responsiveness or not. Interestingly, while four out of the five top ranked proteins are reported to be pH-responsive / regulated, for the bottom five proteins, only one is reported to be regulated by / sensitive to pH (See Supplementary Table 1). This example illustrates how, using patcHwork, it is possible to conduct quantitative pH sensitivity analysis, ranking hundreds of protein sequences in a matter of minutes.

**Figure 1.**
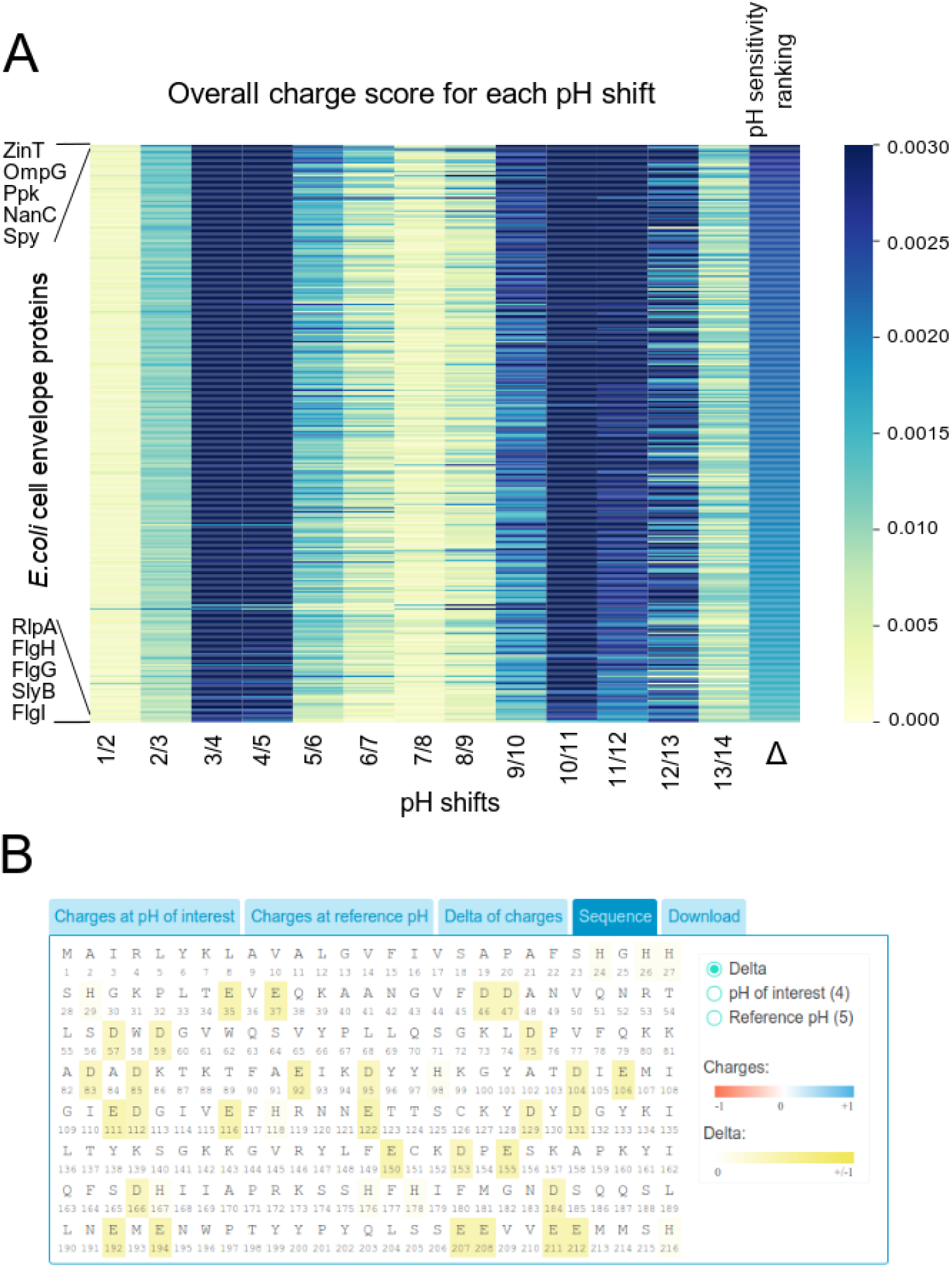
Demonstration of sequence functionality of patcHwork. **A-** Overall charge score based pH sensitivity ranking of 309 *E.coli* cell envelope proteins (GO:0030313) for 1 unit incremental pH increase from pH 1 to 14. Δ represents the mean of overall charge scores for each protein. Note: Uncharacterized proteins are not considered for future literature search given in the Supplementary Table 1. **B-** ZinT, the most pH responsive *E.coli* cell envelope protein from (A) is analyzed for pH shift 4 to 5 with patcHwork by using sequence functionality.

### Structural analysis of pH-induced changes in the sweet taste protein neoculin

To demonstrate how patcHwork can be used to narrow down potential residues likely to be involved in the pH response mechanism of a given protein, we analyzed neoculin, a heterodimeric protein from the plant *Curculigo latifolia*. Neoculin (also called curculin) consists of an acidic (NAS) and a basic (NBS) subunit and it turns sour into sweet taste (32). This effect has been shown to be induced by low pH (33). A mutant neoculin where all five histidines were mutated to alanine is shown to be active across pH conditions (25). Specifically, His11 in the NBS was identified as the main pH sensor responsible for the activity of the protein at low pH. Nakajima and colleagues suggested that low pH-mediated loss of the aromatic interaction between His11 and His14 in the NBS is essential for the pH-responsive function of neoculin (25).

While it is intuitive to mutate histidine residues to investigate the pH response mechanism of a given protein considering that the pKa of histidine is close to the physiological pH, analyzing the protein structure could give further valuable information to narrow candidate residues down as well as give insights into a potential mechanism. For this purpose, we ran a structure-based analysis of neoculin (PDB id: 2D04, chains A and B) (34) with patcHwork for the pH shift from 4 to 8, the two extreme pH values used in the experimental study (25) (Session can be accessed at: https://patchwork.biologie.uni-freiburg.de/results.php?key=example_pdb). We found that the non-covalent bonds between Arg38 - His36 (aromatic interaction), His67 - Ser50 (hydrogen bond), Arg53 - His11 and His11-His14 (aromatic interactions) are destroyed. Interestingly, His11 has two disrupted non-covalent bonds while the other differentially charged residues only have single ones, hinting that a change in pH especially affects this region occurred in the Arg53 - His11 - His14 triad, which is in line with the experimental observations (25) (Figure 2A-B). In addition to the analysis of non-covalent bond changes, we identified a pH-sensitive patch constituted by His11, His14 and His67 (Figure 2C).

**Figure 2.**
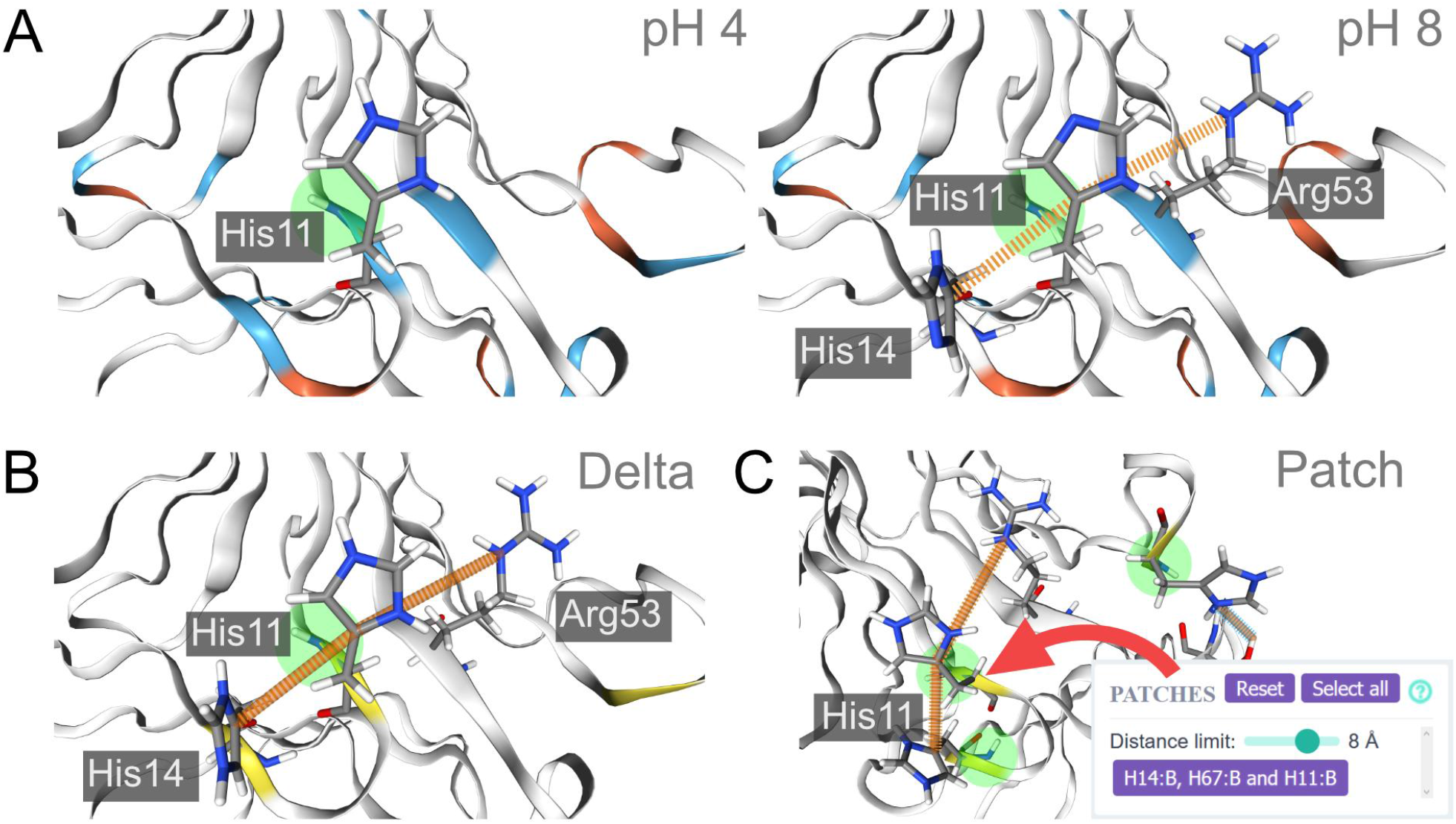
Structure-based patcHwork analysis using neoculin X-ray structure (PDB id: 2D04, chains A and B) for pH 4 and 8. **A-** The intra-molecular pi-pi interaction between His11 and His14 is shown in orange for pH 8. **B**- Delta view in patcHwork. Intra-molecular non-covalent bonds that are present only for one of the two pH values are shown in orange. **C-** Patch view in patcHwork. The pH-sensitive patch, identified when the threshold for the distances between pH-sensitive residues is set ≤ 8 Å, is shown. Individual residues within the patch are displayed in the box. In all panels, the structure of neoculin with both its subunits (PDB ID: 2D04, chains A and B) was used in the analysis.The figure was generated in patcHwork with modifications to the residue labeling. Negatively charged residues are shown in red, positively charged residues in blue and neutral residues in white. Residues that are differentially charged at pH 8 and 4 are highlighted in yellow.

### Structural analysis of pH-regulated mouse anion exchanger 2 (mAE2)

To showcase the usefulness of patcHwork for the pH sensitivity analysis of proteins consisting of over a thousand amino acids, for which the shift in the pKa of residues is due to the surrounding environment, we selected as the last case study an anion exchanger protein (also called bicarbonate transporter).

The anion exchangers 1-3 (AE1-3) mediate Na^+^-independent Cl^-^/HCO_3_^-^ exchange in several types of cells. Their functions range from gas transport, to cell volume and intracellular pH regulation (35). Both AE1 and AE2 catalyze H^+^-SO_4_ ^2-^/Cl^-^ and H^+^-SO_4_^2-^/ H^+^-SO_4_^2-^ exchange in a pH-sensitive fashion: in an acidic environment (pH of 5.5), the H^+^-SO_4_^2-^ efflux is maximal, whereas it sharply decreases at near neutral pH (pH of 7.5) (27–30). In human AE1 (hAE1), the binding site for the proton that is cotransported with SO_4_^2-^ was proposed to be the Glu681(30). Chemically converting this negatively-charged residue to an alcohol (Glu681OH), and thus rendering it neutrally charged, leads to proton independent transport of SO_4_ ^2-^ as well as nearly completely eliminating the transport’s pH dependence (31, 36). Mutating the corresponding conserved glutamic acid in mouse AE1 (mAE1) and AE2 (mAE2), being Glu699 and Glu1007 respectively, to the neutral glutamine abolishes the preference of sulfate exchange in acidic medium similarly to the experiments with hAE1 (27–31, 36).

In the absence of this experimental evidence, we would typically perform mutagenesis analysis on mAE2 to identify amino acids involved in the pH-regulated SO_4_^2-^ transport. For this aim we would likely focus on the histidines, as their pKa are close to the physiological pH and thus we know they would change protonation state for a pH shift from 5.5 to 7.5. This methodology would lead to identifying 35 candidate histidine residues on mAE2, which consists of 1237 amino acids. However, it would not lead to identifying Glu1007, which is reported to be the key pH-responsive amino acid of mAE2 (27–29).

Using patcHwork for the analysis allows avoiding falling in the prototypical histidine-oriented approach. As a matter of fact, by making use of pKa predictions from PROPKA (15), patcHwork is able to consider the surrounding environment of the side chains of each amino acid to calculate the shift in pKa, since this is known to exert a role (12). We downloaded the mAE2 model structure from the AlphaFold Protein Structure Database (37, 38) (Model id: AF-P13808-F1), removing regions with low model confidence score (< 70) with the exception of the linker between the two domains (anion exchanger and cytoplasmic domains), and we submitted the structure to patcHwork using pH of interest of 5.5 and reference pH of 7.5, which were used in the experimental studies (28, 29). We obtain the following three levels of information (Session can be accessed at: https://patchwork.biologie.uni-freiburg.de/results.php?key=mAE2_example): 1-residues that change their protonation state (shown in yellow in patcHwork); 2-residues that change their protonation state and cause non-covalent bond changes (yellow residues with red and green stripes in patcHwork), and 3-patches: residues that change their protonation state and are in close physical proximity with other residues that change their protonation states (green spheres in patcHwork). These three features generated from patcHwork can be used to identify pH-sensitive residues as seen in Table 1.

**Table 1:**
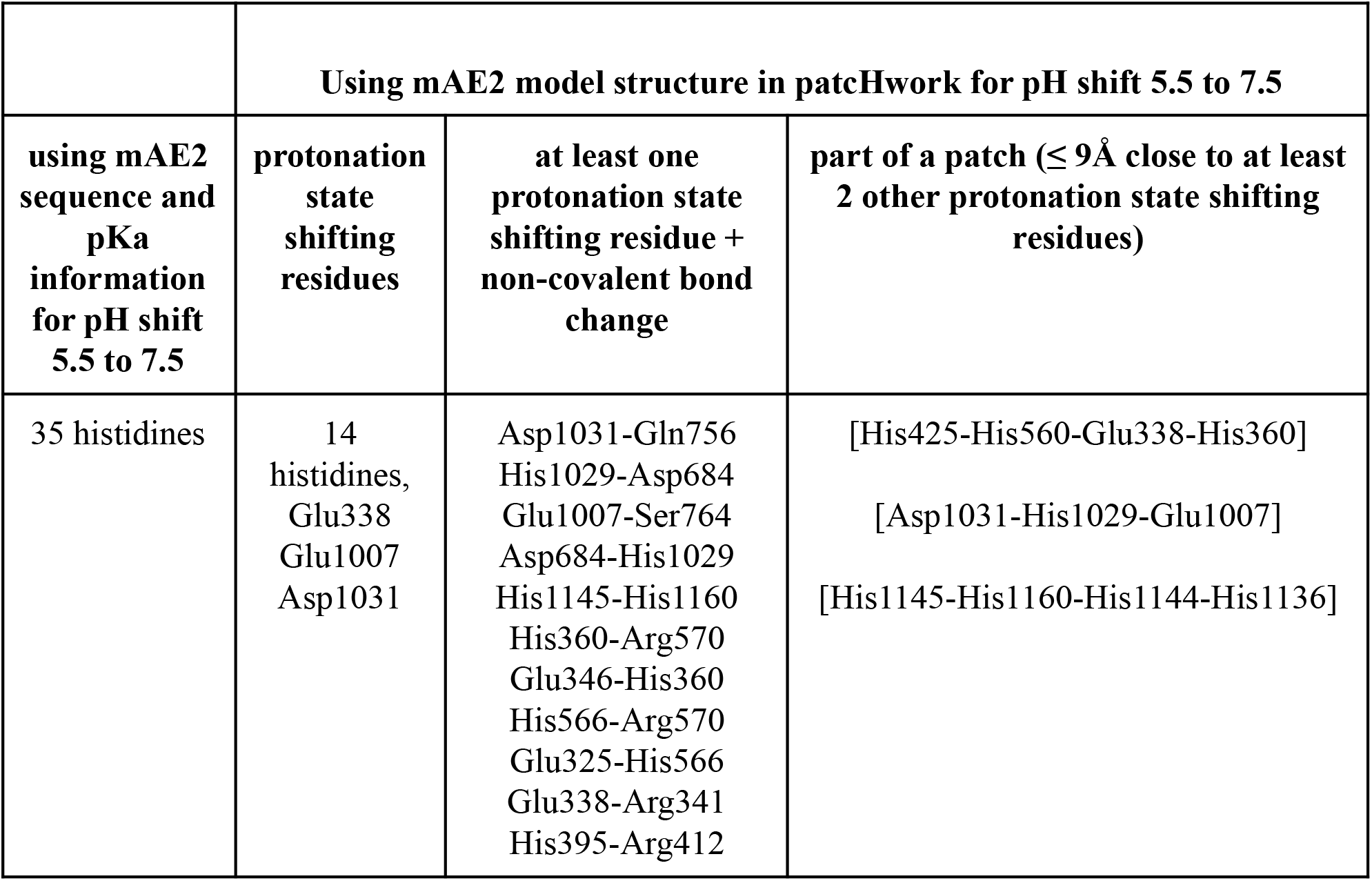
Comparison of histidine-based and structure-based analyses (the latter performed with patcHwork) for the identification of pH-sensitive amino acids within mAE2

|  | Using mAE2 model structure in patchHwork for pH shift 5.5 to 7.5 |  |  |
| --- | --- | --- | --- |
| using mAE2 sequence and pKa information for pH shift 5.5 to 7.5 | protonation state shifting residues | at least one protonation state shifting residue + non-covalent bond change | part of a patch ($\leq 9\text{\AA}$ close to at least 2 other protonation state shifting residues) |
| 35 histidines | 14 histidines, Glu338, Glu1007, Asp1031 | Asp1031-Gln756<br>His1029-Asp684<br>Glu1007-Ser764<br>Asp684-His1029<br>His1145-His1160<br>His360-Arg570<br>Glu346-His360<br>His566-Arg570<br>Glu325-His566<br>Glu338-Arg341<br>His395-Arg412 | [His425-His560-Glu338-His360]<br>[Asp1031-His1029-Glu1007]<br>[His1145-His1160-His1144-His1136] |

Using patcHwork it is possible to group pH-sensitive residues into three patches (Figure 3A), resulting in a broader view approach compared to using histidine-only information with standard pKa values. Looking at the structure, we can speculate that a pH-sensitive region influencing substrate binding is more likely to be found within the anion exchanger domain itself, close to the two substrate binding sites (assuming that these are are conserved with the sites found in hAE1 (38); see Supplementary Information) rather than within the cytoplasmic domain. In this case, the patch constituted by Asp1031 - His1029 - Glu1007 is the most likely pH-sensitive region of mAE2 (Figure 3B). We conclude that, being the closest to the residues involved in substrate binding, residue Glu1007 is the prime candidate for mutational experiments (Figure 3C). Furthermore, as Glu1007 becomes protonated, a shift in the hydrogen bond with Ser764, belonging to the substrate binding pocket, could hint at a restructuring of the binding pocket allowing for a different substrate such as SO_4_^2-^ to bind (Figure 3D). Importantly, our analysis is also in line with an alternative mechanism that was previously proposed, whereby the negatively-charged glutamic acid inhibits sulfate binding through ionic repulsion, which ceases when the glutamic acid is protonated (40).

**Figure 3.**
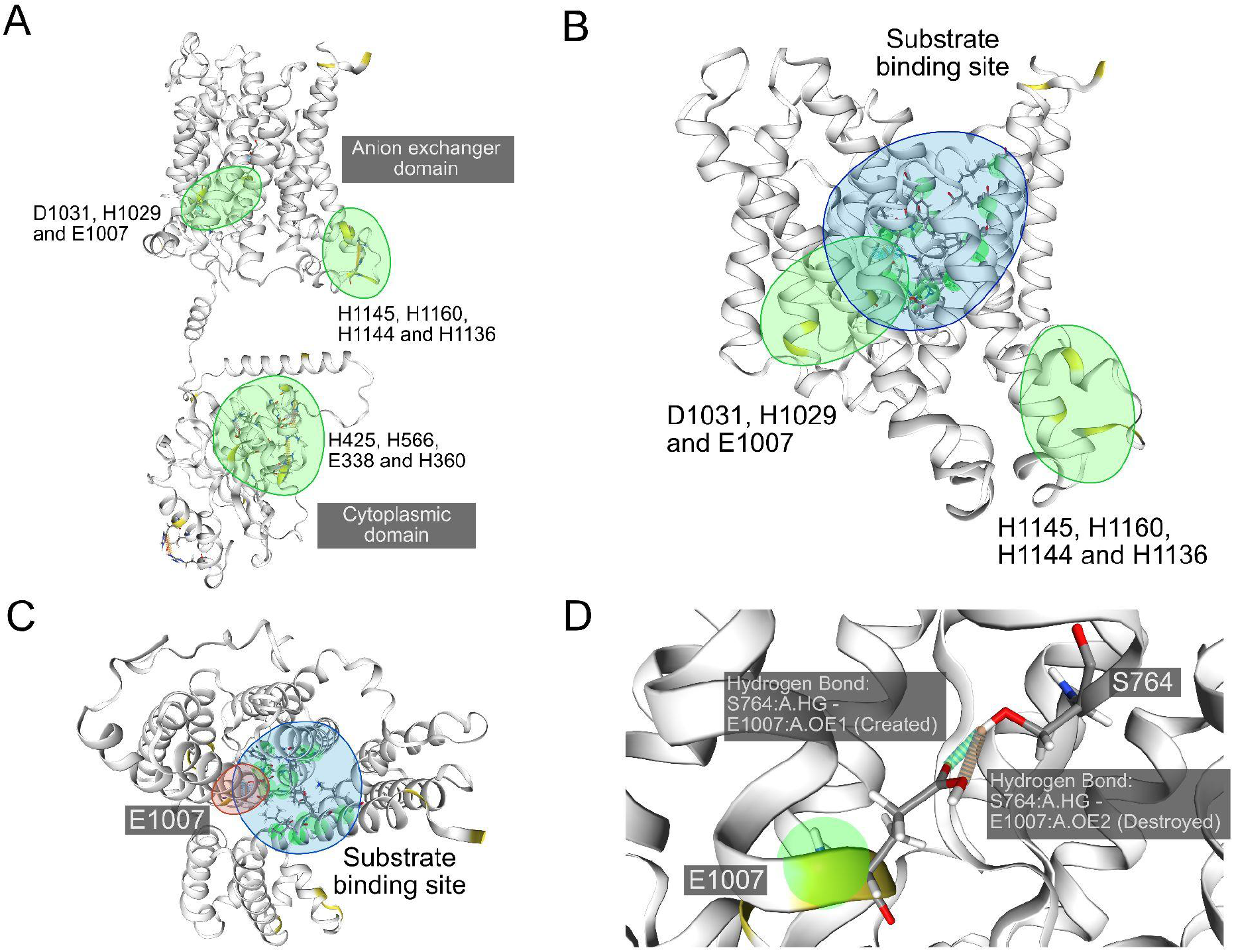
mAE2 structure pH sensitivity analysis by using patcHwork for pH 5.5 and 7.5 **A-** Three respective pH sensitive patches identified are shown in green. **B-** Both patches of the anion exchanger domain surrounded in green are shown with mAE2 substrate binding sites in blue (For details of the residues see Supplementary Information). **C-** Top view of the substrate binding site and differentially charged residue Glu1007 closest to it. **D-** Shifts in the hydrogen bond between Glu1007 and Ser764, the latter being a part of the substrate loop binding site.

## Conclusion

patcHwork is a novel, easy-to-use web server that offers users the possibility to perform high-throughput pH-sensitivity analysis of protein sequences and structures.

A limitation of patcHwork is that it does not capture pH-dependent structural dynamics that can occur upon pH shift. This can be achieved via molecular dynamics simulations at constant pH (41) comparing the results obtained at different pH values. However, such approaches are computationally demanding and require expertise. On a more intuitive and accessible level, patcHwork allows users to nonetheless predict potential structural rearrangements upon evaluation of gain or loss of non-covalent bonds caused by pH shift.

We believe patcHwork will be an invaluable tool supporting research and teaching, deepening our mechanistic understanding of how pH impacts protein function.

## Acknowledgments

We would like to thank members of the Di Ventura laboratory for their valuable feedback and suggestions.

## Data availability

The web-site is freely available at: https://patchwork.biologie.uni-freiburg.de

## References

1. Pace, C.N., Grimsley, G.R. and Scholtz, J.M. (2009) Protein Ionizable Groups: pK Values and Their Contribution to Protein Stability and Solubility. J. Biol. Chem., 284, 13285–13289. https://doi.org/10.1074/jbc.R800080200 http://www.ncbi.nlm.nih.gov/pmc/articles/PMC2679426

2. Schaefer, M., Sommer, M. and Karplus, M. (1997) pH-Dependence of Protein Stability: Absolute Electrostatic Free Energy Differences between Conformations. J. Phys. Chem. B, 101, 1663–1683. https://doi.org/10.1021/jp962972s

3. Mueller, E.A., Iken, A.G., Ali Öztürk, M., Winkle, M., Schmitz, M., Vollmer, W., Di Ventura, B. and Levin, P.A. (2021) The active repertoire of Escherichia coli peptidoglycan amidases varies with physiochemical environment. Mol. Microbiol., 116, 311–328. https://doi.org/10.1111/mmi.14711

4. Pathak, A.K. (2018) Effect of pH on the hinge region of influenza viral protein: a combined constant pH and well-tempered molecular dynamics study. J. Phys. Condens. Matter, 30, 195101. https://doi.org/10.1088/1361-648X/aab98c

5. Zhao, Y.J., He, W.M., Zhao, Z.Y., Li, W.H., Wang, Q.W., Hou,Y.N., Tan, Y. and Zhang, D. (2021) Acidic pH irreversibly activates the signaling enzyme SARM1. FEBS J., 288, 6783–6794. https://doi.org/10.1111/febs.16104 http://www.ncbi.nlm.nih.gov/pubmed/34213829

6. Schönichen, A., Webb, B.A., Jacobson, M.P. and Barber, D.L. (2013) Considering Protonation as a Posttranslational Modification Regulating Protein Structure and Function. Annu. Rev. Biophys., 42, 289–314. https://doi.org/10.1146/annurev-biophys-050511-102349 http://www.ncbi.nlm.nih.gov/pubmed/23451893

7. Karshikoff, A. and Jelesarov, I. (2008) Salt Bridges and Conformational Flexibility: Effect on Protein Stability. Biotechnol. Biotechnol. Equip., 22, 606–611. https://doi.org/10.1080/13102818.2008.10817520

8. Hebditch, M. and Warwicker, J. (2019) Web-based display of protein surface and pH-dependent properties for assessing the developability of biotherapeutics. Sci. Rep., 9, 1969. https://doi.org/10.1038/s41598-018-36950-8

9. Bartonek, L. and Zagrovic, B. (2019) VOLPES: an interactive web-based tool for visualizing and comparing physicochemical properties of biological sequences. Nucleic Acids Res., 47, W632–W635. https://doi.org/10.1093/nar/gkz407

10. Unni, S., Huang, Y., Hanson, R.M., Tobias, M., Krishnan, S., Li, W.W., Nielsen, J.E. and Baker, N.A. (2011) Web servers and services for electrostatics calculations with APBS and PDB2PQR. J. Comput. Chem., 32, 1488–1491. https://doi.org/10.1002/jcc.21720

11. Cameselle, J.C., Ribeiro, J.M. and Sillero, A. (1986) Derivation and use of a formula to calculate the net charge of acid-base compounds. Its application to amino acids, proteins and nucleotides. Biochem. Educ., 14, 131–136. https://doi.org/10.1016/0307-4412(86)90176-7

12. Isom, D.G., Castañeda, C.A., Cannon, B.R. and E,B.G.-M. (2011) Large shifts in pKa values of lysine residues buried inside a protein. Proc. Natl. Acad. Sci., 108, 5260–5265. https://doi.org/10.1073/pnas.1010750108 http://www.ncbi.nlm.nih.gov/pubmed/21389271

13. Alexov, E., Mehler, E.L., Baker, N., Baptista, A.M., Huang,Y., Milletti, F., Nielsen, J.E., Farrell, D., Carstensen, T., Olsson, M.H.M., et al. (2011) Progress in the prediction of pKa values in proteins. Proteins Struct. Funct. Bioinforma., 79, 3260–3275. https://doi.org/10.1002/prot.23189

14. Dolinsky, T.J., Czodrowski, P., Li, H., Nielsen, J.E., Jensen, J.H., Klebe, G. and Baker, N.A. (2007) PDB2PQR: Expanding and upgrading automated preparation of biomolecular structures for molecular simulations. Nucleic Acids Res., 35, W522–5. https://doi.org/10.1093/nar/gkm276 http://www.ncbi.nlm.nih.gov/pubmed/17488841

15. Olsson, M.H.M., Søndergaard, C.R., Rostkowski, M. and Jensen, J.H. (2011) PROPKA3: Consistent Treatment of Internal and Surface Residues in Empirical pKa Predictions. J. Chem. Theory Comput., 7, 525–537. https://doi.org/10.1021/ct100578z

16. Rostkowski, M., Olsson, M.H., Søndergaard, C.R. and Jensen, J.H. (2011) Graphical analysis of pH-dependent properties of proteins predicted using PROPKA. BMC Struct. Biol., 11, 6. https://doi.org/10.1186/1472-6807-11-6 http://www.ncbi.nlm.nih.gov/pmc/articles/PMC3038139

17. Humphrey, W., Dalke, A. and Schulten, K. (1996) VMD: Visual molecular dynamics. J. Mol. Graph., 14, 33–38. https://doi.org/10.1016/0263-7855(96)00018-5 http://www.ncbi.nlm.nih.gov/pubmed/8744570

18. Piovesan, D., Minervini, G. and Tosatto, S.C.E. (2016) The RING 2.0 web server for high quality residue interaction networks. Nucleic Acids Res., 44, W367–W374. https://doi.org/10.1093/nar/gkw315 http://www.ncbi.nlm.nih.gov/pmc/articles/PMC4987896

19. Jubb, H.C., Higueruelo, A.P., Ochoa-Montaño, B., Pitt, W.R., Ascher, D.B. and Blundell, T.L. (2017) Arpeggio: A Web Server for Calculating and Visualising Interatomic Interactions in Protein Structures. J. Mol. Biol., 429, 365–371. https://doi.org/10.1016/j.jmb.2016.12.004 http://www.ncbi.nlm.nih.gov/pmc/articles/PMC5282402

20. Ferruz, N., Schmidt, S. and Höcker, B. (2021) ProteinTools: a toolkit to analyze protein structures. Nucleic Acids Res., 49, W559–W566. https://doi.org/10.1093/nar/gkab375

21. Abraham, M.J., Murtola, T., Schulz, R., Páll, S., Smith, J.C., Hess, B. and Lindahl, E. (2015) GROMACS: High performance molecular simulations through multi-level parallelism from laptops to supercomputers. SoftwareX, 1–2, 19–25. https://doi.org/10.1016/j.softx.2015.06.001

22. Barlow, D.J. and Thornton, J.M. (1983) Ion-pairs in proteins. J. Mol. Biol., 168, 867–885.https://doi.org/10.1016/S0022-2836(83)80079-5

23. Bhattacharyya, R., Samanta, U. and Chakrabarti, P. (2002) Aromatic–aromatic interactions in and around α-helices. Protein Eng. Des. Sel., 15, 91–100. https://doi.org/10.1093/protein/15.2.91

24. Gallivan, J.P. and Dougherty, D.A. (1999) Cation-π interactions in structural biology. Proc. Natl. Acad. Sci. U. S. A., 96, 9459–9464. http://www.ncbi.nlm.nih.gov/pmc/articles/PMC22230

25. Nakajima, K., Yokoyama, K., Koizumi, T., Koizumi, A., Asakura, T., Terada, T., Masuda, K., Ito, K., Shimizu-Ibuka, A., Misaka, T., et al. (2011) Identification and Modulation of the Key Amino Acid Residue Responsible for the pH Sensitivity of Neoculin, a Taste-Modifying Protein. PLOS ONE, 6, e19448. https://doi.org/10.1371/journal.pone.0019448

26. Ohkubo, T., Tamiya, M., Abe, K. and Ishiguro, M. (2015) Structural Basis of pH Dependence of Neoculin, a Sweet Taste-Modifying Protein. PLOS ONE, 10, e0126921. https://doi.org/10.1371/journal.pone.0126921

27. Chernova, M.N., Stewart, A.K., Barry, P.N., Jennings, M.L. and Alper, S.L. (2008) Mouse Ae1 E699Q mediates SO42-i/aniono exchange with [SO42-]i-dependent reversal of wild-type pHo sensitivity. Am. J. Physiol.-Cell Physiol., 295, C302–C312. https://doi.org/10.1152/ajpcell.00109.2008

28. Sekler, I., Lo, R.S. and Kopito, R.R. (1995) A Conserved Glutamate Is Responsible for Ion Selectivity and pH Dependence of the Mammalian Anion Exchangers AE1 and AE2 (*). J. Biol. Chem., 270, 28751–28758. https://doi.org/10.1074/jbc.270.48.28751

29. Reimold, F.R., Stewart, A.K., Stolpe, K., Heneghan, J.F., Shmukler, B.E. and Alper, S.L. (2013) Substitution of transmembrane domain Cys residues alters pHo-sensitive anion transport by AE2/SLC4A2 anion exchanger. Pflüg. Arch. - Eur. J. Physiol., 465, 839–851. https://doi.org/10.1007/s00424-012-1196-6

30. Jennings, M.L. and Smith, J.S. (1992) Anion-proton cotransport through the human red blood cell band 3 protein. Role of glutamate 681. J. Biol. Chem., 267, 13964–13971. https://doi.org/10.1016/S0021-9258(19)49664-6

31. Chernova, M.N., Jiang, L., Crest, M., Hand, M., Vandorpe, D.H., Strange, K. and Alper, S.L. (1997) Electrogenic Sulfate/Chloride Exchange in Xenopus Oocytes Mediated by Murine AE1 E699Q. J. Gen. Physiol., 109, 345–360. http://www.ncbi.nlm.nih.gov/pmc/articles/PMC2217076

32. Shirasuka, Y., Nakajima, K.-I., Asakura, T., Yamashita, H., Yamamoto, A., Hata, S., Nagata, S., Abo, M., Sorimachi, H. and Abe, K. (2004) Neoculin as a new taste-modifying protein occurring in the fruit of Curculigo latifolia. Biosci. Biotechnol. Biochem., 68, 1403–1407. https://doi.org/10.1271/bbb.68.1403 http://www.ncbi.nlm.nih.gov/pubmed/15215616

33. Nakajima, K., Morita, Y., Koizumi, A., Asakura, T., Terada, T., Ito, K., Shimizu-Ibuka, A., Maruyama, J., Kitamoto, K., Misaka, T., et al. (2008) Acid-induced sweetness of neoculin is ascribed to its pH-dependent agonistic-antagonistic interaction with human sweet taste receptor. FASEB J. Off. Publ. Fed. Am. Soc. Exp. Biol., 22, 2323–2330. https://doi.org/10.1096/fj.07-100289 http://www.ncbi.nlm.nih.gov/pubmed/18263698

34. Shimizu-Ibuka, A., Morita, Y., Terada, T., Asakura, T., Nakajima, K., Iwata, S., Misaka, T., Sorimachi, H., Arai, S. and Abe, K. (2006) Crystal Structure of Neoculin: Insights into its Sweetness and Taste-modifying Activity. J. Mol. Biol., 359, 148–158. https://doi.org/10.1016/j.jmb.2006.03.030

35. Romero, M.F., Fulton, C.M. and Boron, W.F. (2004) The SLC4 family of HCO3-transporters. Pflüg. Arch., 447, 495–509. https://doi.org/10.1007/s00424-003-1180-2

36. Jennings, M.L. and Al-Rhaiyel, S. (1988) Modification of a carboxyl group that appears to cross the permeability barrier in the red blood cell anion transporter. J. Gen. Physiol., 92, 161–178. https://doi.org/10.1085/jgp.92.2.161

37. Varadi, M., Anyango, S., Deshpande, M., Nair, S., Natassia, C., Yordanova, G., Yuan, D., Stroe, O., Wood, G., Laydon, A., et al. (2022) AlphaFold Protein Structure Database: massively expanding the structural coverage of protein-sequence space with high-accuracy models. Nucleic Acids Res., 50, D439–D444. https://doi.org/10.1093/nar/gkab1061

38. Jumper, J., Evans, R., Pritzel, A., Green, T., Figurnov, M., Ronneberger, O., Tunyasuvunakool, K., Bates, R., žídek, A., Potapenko, A., et al. (2021) Highly accurate protein structure prediction with AlphaFold. Nature, 596, 583–589. https://doi.org/10.1038/s41586-021-03819-2

39. Zhekova, H.R., Pushkin, A., Kayık, G., Kao, L., Azimov, R., Abuladze, N., Kurtz, D., Damergi, M., Noskov, S.Y. and Kurtz, I. (2021) Identification of multiple substrate binding sites in SLC4 transporters in the outward-facing conformation: Insights into the transport mechanism. J. Biol. Chem., 296, 100724. https://doi.org/10.1016/j.jbc.2021.100724

40. Reithmeier, R.A.F., Casey, J.R., Kalli, A.C., Sansom, M.S.P., Alguel, Y. and Iwata, S. (2016) Band 3, the human red cell chloride/bicarbonate anion exchanger (AE1, SLC4A1), in a structural context. Biochim. Biophys. Acta BBA - Biomembr., 1858, 1507–1532. https://doi.org/10.1016/j.bbamem.2016.03.030

41. Socher, E. and Sticht, H. (2016) Mimicking titration experiments with MD simulations: A protocol for the investigation of pH-dependent effects on proteins. Sci. Rep., 6, 22523. https://doi.org/10.1038/srep22523

