## Supplementary Information for "patcHwork: A user-friendly pH sensitivity analysis web server for protein sequences and structures"

### 1. patcHwork implementation

#### 1.1 Sequence-based pH sensitivity analysis

The amino acid sequence of the protein of interest is uploaded in FASTA format and two pH values are specified (pH of interest and reference pH). Up to ten thousand protein sequences containing the 20 canonical amino acids can be uploaded to patcHwork. If present, non-canonical amino acids are attributed a charge of zero at any pH. The Henderson-Hasselbalch equation (1) (Eq. 1 and 2) is then applied to determine the charge of the side chain of each residue at the two user-specified pH values. Specifically, the charges of the C-terminal carboxyl functional group, as well as of the side chains of Asp, Glu, Cys and Tyr are calculated with the following modified Henderson-Hasselbalch equation (1) (Eq. 1):

$$netZ = \frac{-1}{10^{pKa - pH} + 1} \quad \text{(Eq. 1)}$$

The charges of the N-terminal amino functional group, as well as of the side chains of Arg, His and Lys are calculated with the following modified Henderson-Hasselbalch equation (1) (Eq. 2):

$$netZ = \frac{1}{10^{pH - pKa} + 1} \quad \text{(Eq. 2)}$$

When more than one charged group is present in the side chain of the amino acid, the sum of charges of all groups is calculated.

The difference between net charges of each residue at the two pH values is then calculated (Eq. 3) and given as an output:

$$\Delta Z = netZ_{pHofinterest} - netZ_{referencepH} \quad \text{(Eq. 3)}$$

To allow for the easy comparison of multiple sequences, we provide a normalised pH sensitivity (overall charge score), by dividing the number of pH-sensitive residues by the sequence length. The output is downloadable as a text file.

#### 1.2 Structure-based pH sensitivity analysis

The 3D protein structure is uploaded in PDB format and two pH values are specified (pH of interest and reference pH). Multiple structures (up to 50 structures and maximum 20 MB per .pdb file) can be uploaded at once. The atomic structure is then processed with the PDB2PQR v2.1.1 software (2) with the PARSE forcefield (3) and PROPKA3 (4) options. Importantly, compared to other alternatives, PROPKA was found to be more accurate in predicting pKa values (5) and achieved $<1$ root mean square deviation (RSMD) from experimentally determined pKa values (4). patcHwork runs PDB2PQR twice to generate two structure files (one at the pH of interest and one at the reference pH), which are then parsed to retain only the atomic information of canonical amino acids. For both structure files, each residue's partial atomic charges are summed up to get the residue's total charge. The difference between each residue's charge at the pH of interest and the reference pH is then calculated (Eq. 3).

##### 1.2.1 Intra-molecular non-covalent bond determination on protein structures

A list of proximal pairs of residues –that is, all two residues found at an atomic distance $\leq$ 10 Å from each other– is created at both pH values and the presence of intra-molecular non-covalent bonds between each pair is determined (see below for detailed information). The non-covalent bonds investigated are hydrogen bonds, salt bridges and aromatic (pi-pi and pi-cation) interactions, since these can be affected by the protonation state of the residues. In this way, we are able to visualise the bonds that are created or destroyed when shifting from the reference pH to the pH of interest.

##### 1.2.2 Hydrogen bonds

For hydrogen bonds, we set the angle between the hydrogen donor (D) and acceptor (A) to $\leq 30°$ and the distance between D and A to 3.5 Å as defined in the GROMACS 5.1 molecular dynamics (MD) software package (6). The N, O, S and C atoms can be potential acceptors and/or donors. While they can all be acceptors, atoms can be considered donors only if they have at least one hydrogen atom covalently bound to them. Based on the list of proximal residue pairs, all potential acceptor and donor combinations are evaluated for their compliance to the hydrogen bond criteria. The same residue can form multiple hydrogen bonds and one atom can be involved in hydrogen bonds with multiple residues. A hydrogen bond is considered changed (created or destroyed) when the hydrogen bond definitions are met in one structure but not in the other (pH of interest vs reference pH). This may happen if the orientation and/or the protonation state of residues is/are altered as calculated by PDB2PQR.

##### 1.2.3 Salt bridges

Depending on the pH of the solution, lysine, arginine and histidine residues can be positively charged, whereas aspartic acid, cysteine, glutamic acid and tyrosine can be negatively charged. We consider a salt bridge to be formed between two residues with opposite full electron charges if the distance between any of the side chains' oxygen atoms or the sulphur of cysteine of the negatively charged residue and any of the side chains' nitrogen atoms of the positively charged residue is ≤ 4 Å (7). A salt bridge is created or destroyed if the residue pair satisfies this criterion in one PDB2PQR output structure but not in the other (pH of interest vs reference pH).

##### 1.2.4 Aromatic interactions

Pi-pi interactions are considered established between two aromatic amino acids when only one of the two is positively charged and the centroids of the aromatic rings are not further apart than 7.5 Å (8). Cation-pi interactions are allowed to form between a neutral aromatic amino acid and a positively charged lysine or arginine if the centroid of the aromatic ring and any side chain nitrogen atom of the cation are not further apart than 6 Å (9). If the criteria are met in one PDB2PQR output structure but not in the other (pH of interest vs reference pH), the aromatic interaction is considered created or destroyed.

#### 1.3 pH-sensitive patches

We consider a pH-sensitive patch to be formed if more than two pH-sensitive residues are found within a user-defined distance, set to 8 Å by default. Patches in a given protein or among different proteins can be ranked using the following three scores we calculate and provide in the output file:

Patch score: Number of residues in a patch normalised by the total number of residues in the protein. This score is calculated individually for each patch.

Sum of patch scores: Total number of residues in the patches normalised by the total number of residues in the protein.

Overall charge score: Sum of charges in the protein divided by the total number of residues in the protein.

#### 1.4 Computational resources

patcHwork is divided into two parts: the interactive web server providing the web-based GUI written in PHP and JavaScript and the backend written in C# (.NET 3.0), performing the calculations and generating the downloadable output files. Before structure submission, the user has the option to customise the default intra-molecular non-covalent bond thresholds. After submission, the file(s) is(are) stored for up to 30 days on the server and a unique job key is generated. The file(s) and the user-defined options are then passed to the backend to perform the calculations. While the calculations are being performed, the user can request the current job status. If an email address was provided at submission, a notification will be sent once the job has finished.

patcHworks' backend uses the DotNetZip library (https://github.com/haf/DotNetZip.Semverd) for zipping and unzipping files from the command line or batch files, the MailKit (https://github.com/jstedfast/MailKit) and MimeKit (https://github.com/jstedfast/MimeKit) libraries to email the results to the users. Moreover, as already mentioned, the PDB2PQR v2.1.1 software (2) is used to generate the structure files at the two user-defined pH values.

The web-based GUI uses the the following JavaScript libraries: jQuery (https://github.com/jquery/jquery) and NGL viewer (10) for molecular structure visualisation and figure generation, dygraphs for chart plots (https://github.com/danvk/dygraphs) and vis-network for the bond network visualisation (https://github.com/visjs/vis-network).

### 2. Sequence-based pH sensitivity analysis of *E. coli* cell envelope proteins

947 *E. coli* proteins annotated as being part of the cell envelope (GO:0030313) were identified using the QuickGO web server (11) and downloaded from Uniprot (12). In order to remove duplicates (proteins with sequence similarity over 85%), the FASTA sequences were submitted to the CD-HIT web server (13). The remaining 309 *E. coli* cell envelope protein sequences were submitted to patcHwork and analysed for the pH shifts from 1 to 14 with

increment of 1 (that is, pH of interest 2, reference pH 1; then, pH of interest 3, reference pH 2, and so on). For each pH shift, the overall charge score (total residue charge shift normalised by protein length) was calculated for each protein. Finally, proteins were ranked by their mean overall charge scores for pH shift 1 to 14.

**Table 1:** Five highest and lowest ranked pH-responsive *E. coli* proteins belonging to the cell envelope

| Rank | Uniprot ID | Name | Function | Mechanism |
|---|---|---|---|---|
| Top 1 | P76344 | ZinT | metal binding protein | pH regulated zinc binding (14) |
| Top 2 | P76045 | OmpG | outer membrane protein G | pH regulated pore opening (15) |
| Top 3 | P0A7B1 | Ppk | polyphosphate kinase phosphodiesterase | broad pH activity (16) |
| Top 4 | P69856 | NanC | outer membrane channel protein | ionic strength and pH regulated channel function (17) |
| Top 5 | P77754 | Spy | periplasmic chaperone | - |
| Bottom 5 | P10100 | RlpA | endolytic peptidoglycan transglycosylase | - |
| Bottom 4 | C4ZS19 | FlgH | flagellar L-ring protein | - |
| Bottom 3 | P0ABX5 | FlgG | flagellar basal-body rod protein | pH regulated aggregation (18) |
| Bottom 2 | P0A905 | SlyB | outer membrane lipoprotein | - |
| Bottom 1 | P0A6S3 | FlgI | flagellar P-ring protein | - |

#### 3. Identification of mAE2 substrate bindings sites

Two substrate binding sites in human AE1 were previously identified in molecular dynamics simulations: The first binding site (entry binding site) being composed of the following residues P419, F423, F464-G466, I528, I531, F532, E535, K539, T727, V729, R730, K851. The second binding site (central binding site) is composed of the residues: I528, F532, T727-S731 (19). For mAE2, the homologous residues are defined as P718, F722, F763-G765, I827, I830, F830, E834, K838, A1053, V1055, R1056, M1177 for the entry binding site and I827, F831, A1053-S1057 for the central binding site, the sequence alignment was done with Clustal Omega (20). Residues belonging to both binding sites are shown with green spheres and are all encircled in blue in Figure 3B-C .

**References:**


1. Cameselle,J.C., Ribeiro,J.M. and Sillero,A. (1986) Derivation and use of a formula to calculate the net charge of acid-base compounds. Its application to amino acids, proteins and nucleotides. *Biochem. Educ.*, 14, 131–136.

https://doi.org/10.1016/0307-4412(86)90176-7

2. Dolinsky,T.J., Czodrowski,P., Li,H., Nielsen,J.E., Jensen,J.H., Klebe,G. and Baker,N.A. (2007) PDB2PQR: Expanding and upgrading automated preparation of biomolecular structures for molecular simulations. *Nucleic Acids Res.*, 35, W522-5.

https://doi.org/10.1093/nar/gkm276

http://www.ncbi.nlm.nih.gov/pubmed/17488841

3. Sitkoff,D., Sharp,K.A. and Honig,B. (1994) Accurate Calculation of Hydration Free Energies Using Macroscopic Solvent Models. *J. Phys. Chem.*, 98, 1978–1988.

https://doi.org/10.1021/j100058a043

4. Olsson,M.H.M., Søndergaard,C.R., Rostkowski,M. and Jensen,J.H. (2011) PROPKA3: Consistent Treatment of Internal and Surface Residues in Empirical pKa Predictions. *J. Chem. Theory Comput.*, 7, 525–537.

https://doi.org/10.1021/ct100578z

5. Davies,M.N., Toseland,C.P., Moss,D.S. and Flower,D.R. (2006) Benchmarking pKa prediction. *BMC Biochem.*, 7, 18.

https://doi.org/10.1186/1471-2091-7-18

6. Abraham,M.J., Murtola,T., Schulz,R., Páll,S., Smith,J.C., Hess,B. and Lindahl,E. (2015) GROMACS: High performance molecular simulations through multi-level parallelism from laptops to supercomputers. *SoftwareX*, 1–2, 19–25.

https://doi.org/10.1016/j.softx.2015.06.001

7. Barlow,D.J. and Thornton,J.M. (1983) Ion-pairs in proteins. *J. Mol. Biol.*, 168, 867–885.

https://doi.org/10.1016/S0022-2836(83)80079-5

8. Bhattacharyya,R., Samanta,U. and Chakrabarti,P. (2002) Aromatic–aromatic interactions in and around α-helices. *Protein Eng. Des. Sel.*, 15, 91–100.

https://doi.org/10.1093/protein/15.2.91

9. Gallivan,J.P. and Dougherty,D.A. (1999) Cation-π interactions in structural biology. *Proc. Natl. Acad. Sci. U. S. A.*, 96, 9459–9464.

http://www.ncbi.nlm.nih.gov/pmc/articles/PMC22230

10. Rose,A.S., Bradley,A.R., Valasatava,Y., Duarte,J.M., Prlić,A. and Rose,P.W. (2018) NGL viewer: web-based molecular graphics for large complexes. *Bioinformatics*, 34, 3755–3758.

https://doi.org/10.1093/bioinformatics/bty419

11. Binns,D., Dimmer,E., Huntley,R., Barrell,D., O'Donovan,C. and Apweiler,R. (2009) QuickGO: a web-based tool for Gene Ontology searching. *Bioinformatics*, 25, 3045–3046.

https://doi.org/10.1093/bioinformatics/btp536

12. The UniProt Consortium (2021) UniProt: the universal protein knowledgebase in

2021. *Nucleic Acids Res.*, 49, D480–D489.

https://doi.org/10.1093/nar/gkaa1100

13. Fu,L., Niu,B., Zhu,Z., Wu,S. and Li,W. (2012) CD-HIT: accelerated for clustering the next-generation sequencing data. *Bioinforma. Oxf. Engl.*, 28, 3150–3152.

https://doi.org/10.1093/bioinformatics/bts565

http://www.ncbi.nlm.nih.gov/pmc/articles/PMC3516142

14. Bellotti,D., Rowińska-Żyrek,M. and Remelli,M. (2020) Novel insights into the metal binding ability of ZinT periplasmic protein from Escherichia coli and Salmonella enterica. *Dalton Trans.*, 49, 9393–9403.

https://doi.org/10.1039/D0DT01626H

15. Structure of the monomeric outer‐membrane porin OmpG in the open and closed conformation | The EMBO Journal.

16. Haeusler,P.A., Dieter,L., Rittle,K.J., Shepler,L.S., Paszkowski,A.L. and Moe,O.A. (1992) Catalytic properties of Escherichia coli polyphosphate kinase: an enzyme for ATP regeneration. *Biotechnol. Appl. Biochem.*, 15, 125–133.

http://www.ncbi.nlm.nih.gov/pubmed/1316760

17. Giri,J., Tang,J.M., Wirth,C., Peneff,C.M. and Eisenberg,B. (2012) Single-channel measurements of an N-acetylneuraminic acid-inducible outer membrane channel in Escherichia coli. *Eur. Biophys. J.*, 41, 259–271.

https://doi.org/10.1007/s00249-011-0781-5

18. Saijo-Hamano,Y., Uchida,N., Namba,K. and Oosawa,K. (2004) In vitro characterization of FlgB, FlgC, FlgF, FlgG, and FliE, flagellar basal body proteins of Salmonella. *J. Mol. Biol.*, 339, 423–435.

https://doi.org/10.1016/j.jmb.2004.03.070

http://www.ncbi.nlm.nih.gov/pubmed/15136044

19. Zhekova,H.R., Pushkin,A., Kayık,G., Kao,L., Azimov,R., Abuladze,N., Kurtz,D., Damergi,M., Noskov,S.Y. and Kurtz,I. (2021) Identification of multiple substrate binding sites in SLC4 transporters in the outward-facing conformation: Insights into

the transport mechanism. *J. Biol. Chem.*, 296, 100724.

https://doi.org/10.1016/j.jbc.2021.100724

20. Sievers,F., Wilm,A., Dineen,D., Gibson,T.J., Karplus,K., Li,W., Lopez,R., McWilliam,H., Remmert,M., Söding,J., *et al.* (2011) Fast, scalable generation of high-quality protein multiple sequence alignments using Clustal Omega. *Mol. Syst. Biol.*, 7, 539.

https://doi.org/10.1038/msb.2011.75

http://www.ncbi.nlm.nih.gov/pmc/articles/PMC3261699
